# Errors in action timing and inhibition facilitate learning by tuning distinct mechanisms in the underlying decision process

**DOI:** 10.1101/153676

**Authors:** Kyle Dunovan, Timothy Verstynen

## Abstract

Goal-directed behavior requires integrating action selection processes with learning systems that adapt control using environmental feedback. These functions intersect in the basal ganglia (BG), which has at least two targets of plasticity: a dopaminergic modulation of striatal pathways and cortical modulation of the subthalamic nucleus (STN). Dual learning mechanisms suggests that feedback signals have a multifaceted impact on BG-dependent decisions. Using a hybrid of accumulation-to-bound decision models and reinforcement learning, we modeled the performance of humans in a stop-signal task where participants (N=75) learned the prior distribution of the timing of a stop signal through trial-and-error feedback. Changes in the drift-rate of the action execution process were driven by errors in action timing, whereas adaptation in the boundary height served to increase caution following failed stops. These findings highlight two interactive learning mechanisms for adapting the control of goal-directed actions based on dissociable dimensions of feedback error.

**Author Summary:** Many complex behavioral goals rely on one’s ability to regulate the timing of action execution while also maintaining enough control to cancel actions in response to “Stop” cues in the environment. Here we examined how these two fundamental components of behavior become tuned to the control demands of the environment by combining principles of reinforcement learning with accumulator models of decision making. The synthesis of these two theoretical frameworks is motivated by previous work showing that reinforcement learning and control rely on overlapping circuitry in the basal ganglia. Leveraging knowledge about the interaction of learning and control signals in this network, we formulated a computational model in which performance feedback is used to modulate key mechanisms of the decision process to facilitate goal acquisition. Model-based analysis of behavioral data collected on an adaptive stop-signal task revealed two critical learning mechanisms: one that adjusts the accumulation rate of the “Go” signal to errors in action timing and another that exercises caution by raising the height of the execution boundary after a failed Stop trial. We show how these independent learning mechanisms interact over the course of learning, shedding light on the behavioral effects plasticity in different pathways of the basal ganglia.

## Introduction

Environmental uncertainty demands that goal-directed actions be executed with a certain degree of caution, requiring agents to strike the appropriate balance between speed and control based on internal goals and contextual constraints. Because of the pervasive and dynamic nature of uncertainty in the real world, the precise amount of behavioral control is rarely a known quantity, and must therefore be learned through trial-and-error. Despite the underlying intuition that intelligent behavior requires both the ability to control actions as well as to learn from consequent feedback, it is not well understood how these learning and control systems interact. One reason for this is that computational models of learning [1,2] and control [3–5] have historically emerged from disparate lines of empirical research (see [6,7] for exceptions), adding difficulty to the already challenging task of inferring cognitive phenomena from gross behavioral measures. Recently, however, insights from cognitive and computational neuroscience have begun to shed light on the interaction of cognitive processes in neural circuits, providing additional empirical anchors for grounding theoretical assumptions [8–11].

The basal ganglia (BG), a subcortical network that shares dense, reciprocal connections with much of cortex, as well as many subcortical neuromodulators, is known to play a critical role in both learning and control processes - acting as a central integration hub where cortically-distributed control commands [12–14], sensory evidence [15,16], and decision variables [17–19] can be weighed and synthesized with feedback-dependent learning signals to facilitate goal-directed behavior [7,20,21]. Cortical information enters the BG through three primary pathways: the first two of these pathways, entering via the striatum, are the direct (i.e., facilitating; Fig 1A, green) and indirect pathways (i.e., suppressing; Fig 1A, blue). The third, hyperdirect (i.e., braking; Fig. 1A, red) pathway, enters via the subthalamic nucleus (STN). Given the various sites of structural overlap between the direct and indirect pathways, including their convergence in the output of the BG [22], we recently proposed that, rather than acting as independent “go” and “no-go” levers, these pathways engage in dynamic competition for control over BG-output [23]. From this perspective, the direct and indirect pathway competition over action execution can be thought of as debate between a “believer” and a “skeptic” (Fig. 1B): activation of the direct pathway increases with evidence supporting action execution and gradually suppresses skepticism of the indirect pathway until a threshold is reached and the action can be gated (Fig. 1B). In contrast, increasing activation of the indirect pathway can proactively suppress or delay action execution in the context of high uncertainty, simultaneously affording more time to reactively cancel action cancellation in response to external stop cues that are mediated by signals from the hyperdirect pathway [13,24].

**Fig 1.**
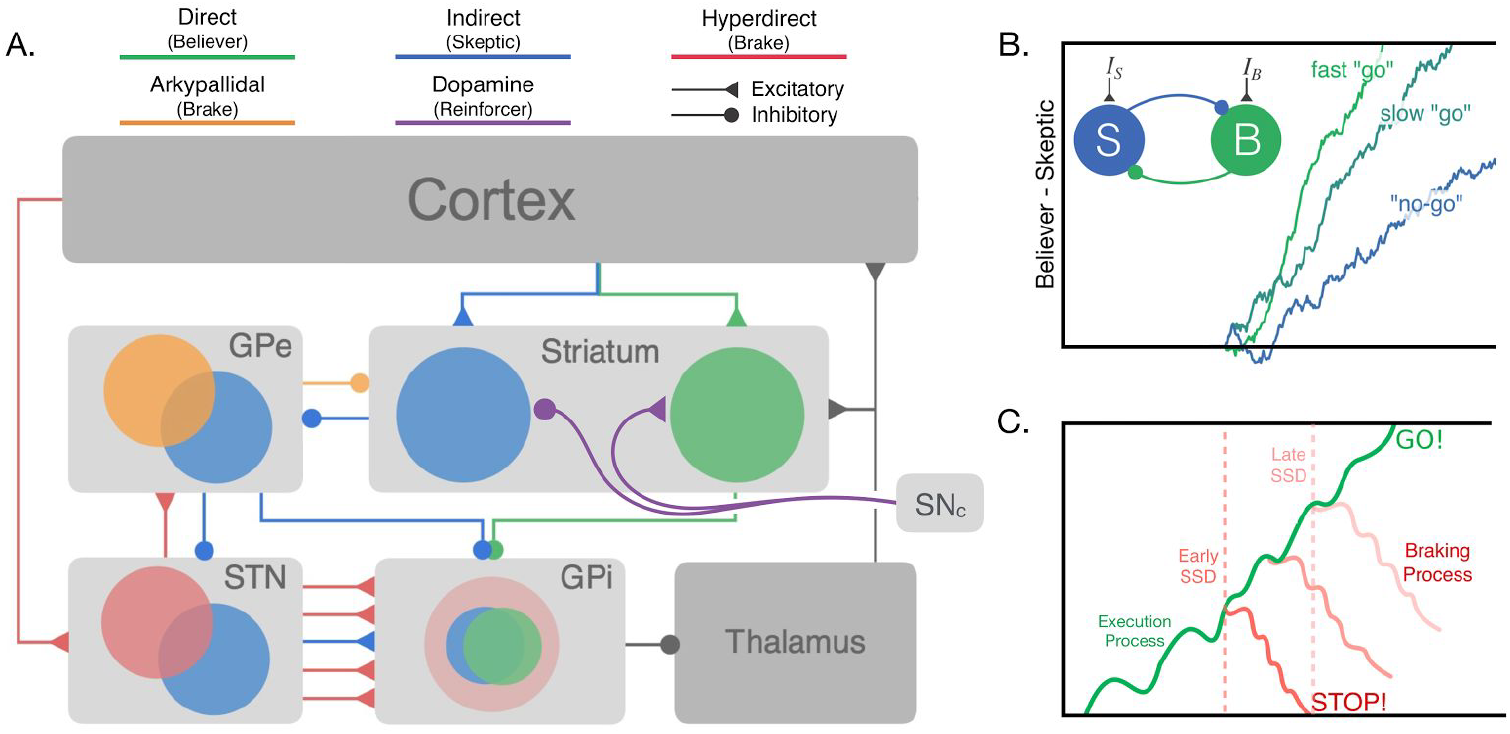
Basal ganglia circuitry and dependent process model. **(A)** Major cortico-basal ganglia (BG) pathways and hypothesized control functions. Projections from the substantia nigra (SNc) deliver dopaminergic feedback signals to the striatum that have opposing effects on the excitability of direct and indirect pathway neurons. **(B)** Competition between direct and indirect pathways represented by mutually inhibiting “believer” (green; direct pathway) and “skeptic” (blue; indirect pathway) populations. Circuit-level dynamics of this competition modulate the rate of evidence accumulation leading up to action execution, leading to faster actions when competition is dominated by “believer”. **(C)** The Dependent Process Model (DPM) assumes that the state of an accumulating execution process at the time a stop cue is registered determines initial state of the braking process, making it more difficult to cancel actions closer to the execution boundary. Panels **A** and **B** have been adapted with permission from Dunovan and Verstynen [23]. Panel **C** has been adapted with permission from Dunovan et al., [13].

A reasonable corollary of the observed convergence of the direct, indirect, and hyperdirect pathways in the output of the BG is that proactive speeding of execution decisions should also influence the efficacy at which the hyperdirect pathway is able to brake the planned execution. Likewise, proactively slowing the accumulation of “go” activity should not only slow the timing of the execution, but also support easier reactive inhibition by reducing the buildup of “go” activity that must be overcome by the hyperdirect pathway in order to brake the action. To test this intuition, we proposed a variant of the classic independent race model in which reactive braking signals, delivered through the hyperdirect pathway, are assumed to be functionally dependent on the current state of the competition between the direct and indirect pathways. In this dependent process model (Fig. 1C) the drift-rate (*v*_E_) of the execution process captures the relative activation of “believer” and “skeptic” populations, leading to a “go“ response when the execution process crosses the upper threshold (*a*), reflecting combined strength of motor-inhibiting forces on basal-ganglia output (e.g., hyperdirect pathway, basal firing rate of BG output nucleus). In the event of a stop-cue, a second braking process is instantiated at the current state of the execution process and must reach the bottom boundary before the execution threshold is reached in order to cancel motor output. This nested dependency of the braking process on the accumulating execution process imposes a speed-control tradeoff, such that increasing the execution drift-rate has direct negative consequences on control, making it more difficult for the braking process to override and cancel action output. While this dependent process model was able to account for the relationship between proactive control and reactive stopping ability, as well as predict BG output during proactive control [13], it fails to account for situations in which the control demands are unknown and thus, must be learned from environmental feedback.

Converging lines of evidence from experimental optogenetics [25] and advances in computational modeling [24] suggest that both primary input structures of the BG, the striatum and STN, are critical for guiding adaptive behavior. For example, similar to classic theories of action-value learning in the striatum, Yttri et al. [25] found that optogenetic reinforcement of cortical input to direct and indirect pathways led to incremental changes in movement velocity, and that this effect disappeared in the presence of dopamine antagonist. Outside the striatum, multiple lines of evidence now point to the STN as a major source of behavioral adaptation in the BG [9,17,26,27]. In line with the notion of parallel learning systems, Wei and Wang [24] showed in a spiking model of the cortico-BG circuitry that plasticity in the striatal and subthalamic pathways gives rise to feedback dependent changes in inhibitory control. Consistent with the dependent process model proposed in Dunovan et al., [13], increasing the connectivity strength of striatal projections in this network model both increased the speed of the execution decision and delayed the stop-signal reaction time. Indeed, this observation lends biological support to the assumptions displayed in Fig 1C about the nested nature of proactive and reactive control signals within the BG. More importantly, however, it suggests distinct functional roles for the two plasticity systems in the BG. First, plasticity in the striatum contributes to adaptive inhibitory control by modulating the drift-rate of the execution process. The STN, on the other hand, should exert control on the height of the execution boundary, delaying action execution in contexts of high uncertainty [17]. Indeed, Cavanaugh et al., [26] found that activity in the STN tracked the degree to which subjects slowed responding after committing an error and that this behavioral phenomenon was described by a diffusion model in which errors led to an increase in threshold on subsequent trials. Taken together, these two BG-dependent learning mechanisms predict two means of adapting action control to environmental feedback: a mechanism for optimizing the timing, or vigor, of action execution as well as a post-error slowing mechanism to impose caution on future action decisions.

In order to test the predictions of this dual mechanism hypothesis, we modeled performance of human participants in an adaptive version of the stop-signal task. Specifically, we show how a straightforward hybridization of the dependent process model [13,23] with principles of reinforcement learning [1], captures feedback-dependent changes in RT and stop accuracy through adaptive changes in the execution drift-rate and boundary height. Drawing on wide ranging evidence from previous studies [9,13,15–17,23–26,28,29], we argue that these behaviorally derived signatures of learning are consistent with dopaminergic tuning of corticostriatal pathways (e.g., drift-rate adaptation) and error-related changes in corticosubthalamic pathways (e.g., boundary height adaptation).

## Results

### Inhibitory control adapts to contextual statistics

Subjects performed an anticipatory version of the stop-signal task (see Fig 2A) similar to that reported previously [13]. On Go trials (n=600), subjects were instructed to make a button press as soon as a vertically rising bar intersected a line near the top of the screen (always occurring at 520 ms) and received a feedback score telling them how early or late their response was (max 100+ pts). On Stop trials, the bar stopped and turned red before reaching the line, signaling the subject to withhold their response. On Context Stop trials (n=200) the stop-signal delay (SSD), defined as the time between trial onset and initiation of the stop-signal, was sampled from one of three probability distributions (Fig. 2B) depending on which group the subject was assigned to. In the Uniform context group, the SSD was sampled uniformly between 10 and 510 ms. In the Early and Late context groups, the SSD was sampled from a normal distribution with means at 250 ms and 350 ms, respectively, both with a standard deviation of 35 ms. To evaluate the effects of Context SSDs on stopping accuracy, Probe Stop trials (n=80) were randomly interspersed throughout all trial blocks with SSDs fixed at 200, 250, 300, 350, or 450 ms (16 trials each). In this paradigm, the Context SSDs establish a prior distribution on the probability of the stop-signal timing, which is known to influence the dynamics of inhibitory control [30]. Shifting the mean of the prior distribution of SSDs later in the trial introduces a greater degree of conflict between Go and Stop trial goals. For instance, the majority of stop-signals in Late context occur too late in the trial to effectively cancel a movement intended to occur at 520 ms, requiring subjects to delay their responding on Go trials to achieve the same level of accuracy as subjects in the Early context.

**Fig 2.**
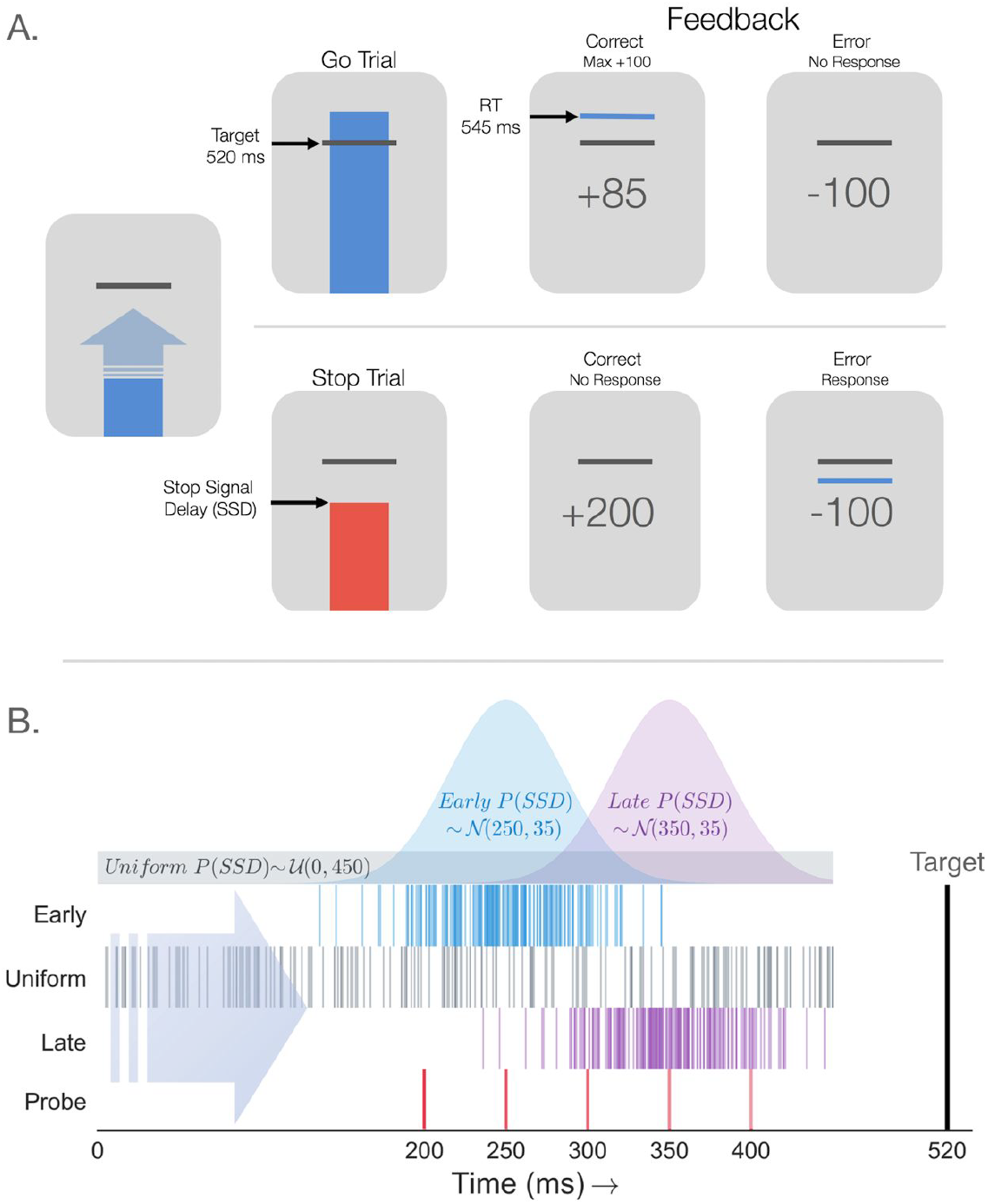
Adaptive stop-signal task and contextual SSD statistics. **(A)** Anticipatory stop-signal task. On Go trials (upper) subjects were instructed to press a key when the ascending bar crossed a target line, always occurring on 520 ms after trial onset. Feedback was given informing the subject if their response was earlier or later than the Go Target (max +100 points). On Stop trials (lower), the bar stopped and turned red prior to reaching the Target line. If no response was made (correct), the subject received a bonus of +200 points. Failure to inhibit the keypress resulted in a −100 point penalty. **(B)** Stop-Signal statistics across Contexts. Distributions show the sampling distributions for SSDs on Context trials in the Early (blue), Uniform (gray), and Late (purple) groups. Early and Late SSDs were Normally distributed (parameters stated as in-figure text 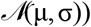. Below the distributions, each row of tick-marks shows the Context SSDs for a single example subject each group. Bottom row of red tick-marks shows the five Probe SSDs included for all subjects regardless of context.

To assess behavioral differences across contexts, we compared accuracy on stop-signal trials at each Probe SSD across groups as well as the mean RTs on correct (response on Go trial) and error (i.e., response on Stop trial) responses. Separate one-way ANOVAs revealed a significant main effect of context across groups on both correct RTs, *F*(2,72)=10.07, *p*<.001, and error RT (responses on stop-signal trials), *F*(2,72)=21.72, *p*<.00001. Consistent with our hypothesis, we found a significant interaction between context condition and Probe SSD, *F*(2.23,80.15)=3.60, *p*=.027 (Fig. 3A). Shifting the mean of the prior distribution on the stop signal later into the trial led to delayed responding on Go trials as well as greater stopping success across Probe trial SSD’s, exhibited by the rightward shift in the stop-curve and RT distributions in the Uniform and Late groups relative to the Early group. Thus, as predicted, participants can reliably learn to modulate their inhibitory control efficiency based on the probabilistic structure of prior stop-signal timing (see also [30]).

**Fig 3.**
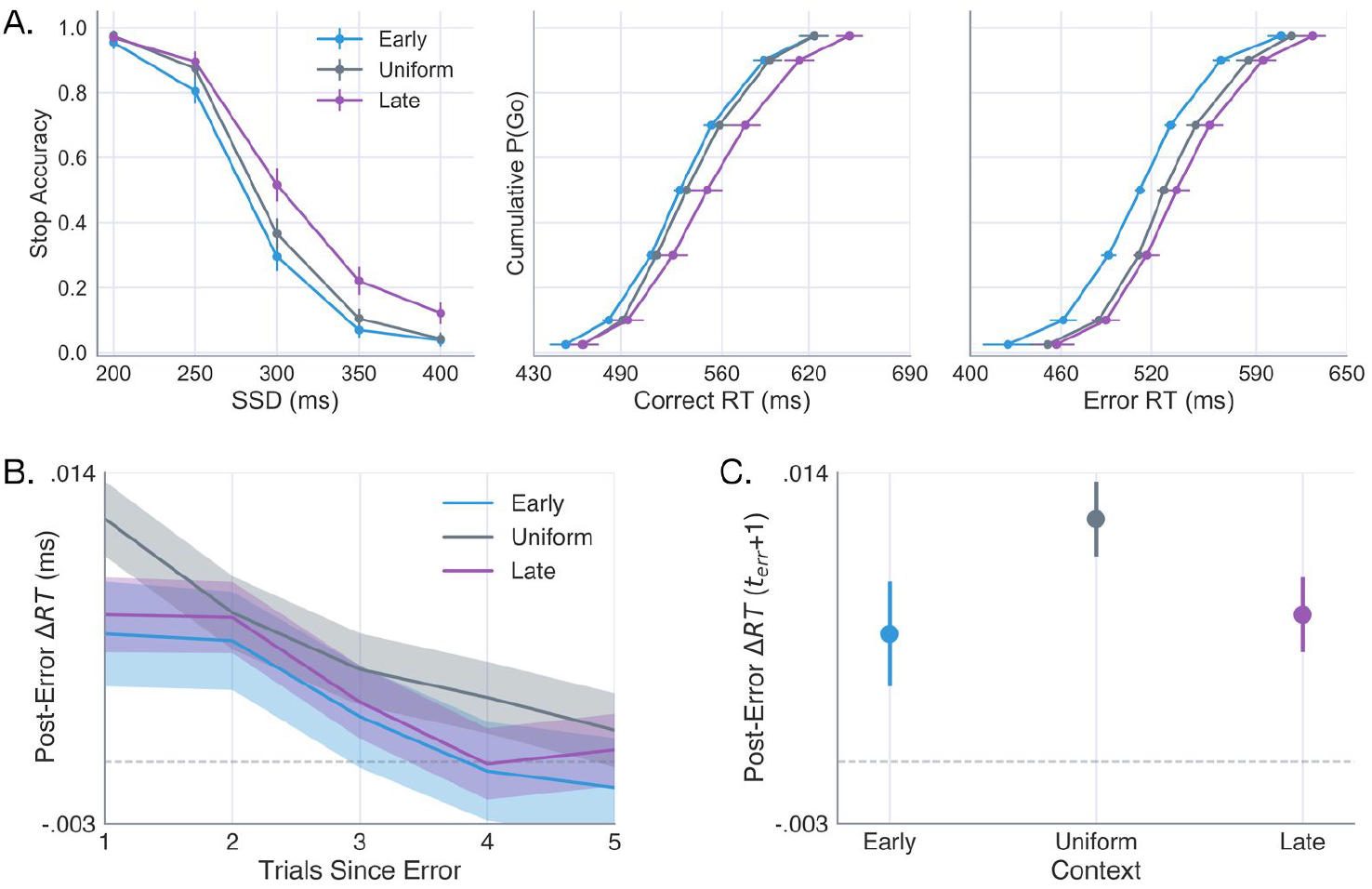
Effects of context on stop accuracy and response times. (A) Subject-averaged stop-accuracy (left) and cumulative RT distributions for correct (Go trials; middle) and error (Stop trials; right) responses in the Early (blue), Uniform (gray), and Late (purple) Contexts. **(B)** Post-error slowing following failed Stop trials in each context and subsequent decay over five trials **(C)** The post-error slowing observed immediately after a failed stop (*t*_*err*_+1) in each Context (e.g., first data point in panel **B**). Error bars and shaded area reflect the 95% confidence interval (CI) calculated across subjects.

We next examined whether failed stop trials elicited any systematic changes in RT on subsequent trials. Fig 3B shows the immediate slowing and subsequent decay in RTs following a stop error, calculated with respect to the average RT on the ten trials that preceded the error. A one-way ANOVA revealed a significant effect of Context on the degree to which subjects slowed responses immediately following stop errors, *F*(2,72)=4.27, *p*=.018. Unlike the observed effects on RT and accuracy, which scaled with differences in the mean SSD in each Context, group differences in post-error slowing appeared to be driven by the variance of SSDs, with stop errors eliciting greater slowing in the Uniform context than in the Early and Late contexts (Fig. 3C). Collectively, these findings suggest that adaptive control is sensitive to multiple task dimensions and that these dimensions manifest in dissociable behavioral profiles.

### Dependent process model of inhibitory control

The dependent process model [31] assumes that an execution decision is made when an accumulating execution process, with onset-time *tr* and drift-rate *v*_E_, crosses an upper decision boundary *a*. The execution process is also influenced by an urgency signal which increases the probability that a response will be executed over time in proportion to a gain parameter *γ* (see Computational Models section of Methods). On Stop trials, a nested braking process, with negative drift-rate *v*_B_, is initiated at the current state of the execution process at the time of the SSD and accumulates back towards the lower boundary (always set equal to 0; see Fig. 1C). The model successfully cancels an action when the braking process reaches the lower bound before execution process terminates at the upper execution threshold.

For a cognitive model to be informative, it is important to verify that its parameters are identifiable, or able to be reliably estimated from observable measures of the target behavior. The issue of model identifiability is particularly relevant to novel variants of sequential sampling models, as several recently proposed models within this class have been found to exhibit poor identifiability despite providing convincing fits to experimental data [32,33]. More common variants, however, such as the drift-diffusion model (DDM) and linear ballistic accumulator, are reasonably identifiable with sufficient trial counts and the application of appropriate optimization procedures [34–36]. In practice, the identifiability of a model can be assessed by performing fits to simulated data, for which the true parameters are known, and comparing the recovered estimates. To evaluate the identifiability of parameters in the DPM we adopted the following procedure. First, we identified three generative parameter sets that approximated the average stopping accuracy curve and RT distributions observed in each Context condition, ensuring that the the generative parameter sets yielded plausible behavioral patterns. Each of these three generative parameter sets served as hyperparameters describing the mean of a normally distributed population from which twenty-five “subject-level” parameter sets were sampled and used to simulate 1000 trials (see Methods for sampling details). This produced a simulated dataset similar in size and dimension to that of the empirical data while capturing the assumption that subject parameter values in each Context vary around a shared mean. Each of the three group-level parameter sets was used to generate 20 simulated datasets (each comprised of 25 randomly sampled subjects with 1000 trials pers subject). The DPM was then fit to the subject-averaged stop-accuracy and RT quantiles for each of the simulated datasets following the optimization routine outlined in the Methods (i.e., here, “subject-averaged” data was calculated by first estimating the stop accuracy curve over probe SSDs, correct RT quantiles, and error RT quantiles for each subject then calculating the mean for each of these values across subjects).

Parameter estimates recovered from the fits are summarized in Fig 4A, with the recovered values for each parameter plotted against the respective generative value for each of the three sets. All parameters were recovered with a high degree of accuracy, with the exception of the urgency gain parameter which was estimated within close range of the generative values but failed to capture mean differences between the three sets. In addition to accurately recovering most generative parameter values, the DPM provided high quality fits to the datasets generated from all three parameter sets, as shown by the positively skewed distribution of *χ*^2^ values in Fig 4A (lower-right) and overlap between observed and predicted stop accuracy and RT quantiles in Fig 4B. The results of this simulation and recovery analysis suggest that parameters of the DPM are identifiable when fitting group-level data and are robust to variability in the parameter values of individual subjects.

**Fig 4.**
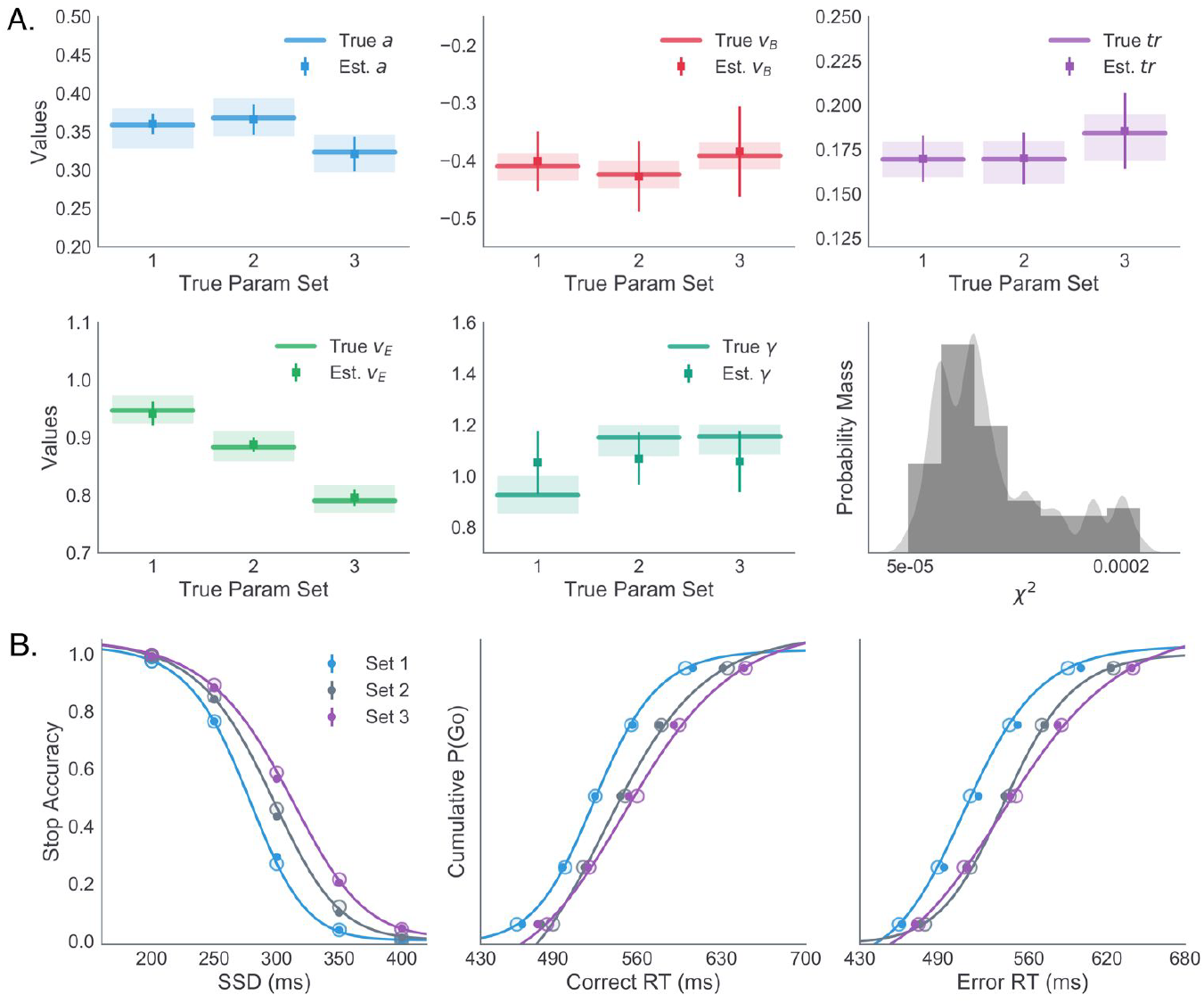
Simulation and parameter recovery analysis of DPM. **(A)** True and estimated boundary height (*a*; top-left, blue), braking drift-rate (*v*_B_; top-middle, red), onset-time (*tr*; top-right, purple), execution drift-rate (*v*_E_; bottom-left, green), and urgency gain (γ; bottom-middle) for three generative parameter sets. True generative parameter means are plotted as lines with the range of sampled subject-level estimates shown in lighter colors. Estimated parameter means are plotted as square markers with error bars reflecting +/- 1 standard deviation. Distribution of χ^2^ values for fits to all 60 simulated datasets is shown in the lower-left panel (gray). (**B)** Stop accuracy curve (left) and correct (middle) and error (right) RT quantiles simulated from true parameter sets 1 (blue), 2 (gray), and 3 (purple) shown as filled circles with average model-predicted values shown as lines and transparent circles.

### Contextual modulation of decision process

We next set out to identify which components of the decision process are adjusted by trial-and-error feedback as participants learn the prior probability of stop signal timing in their assigned context. In order to narrow the search space for feedback-sensitive parameters, we performed the following model fit analyses. First, in order to identify a baseline set of parameters in the DPM, parameters were optimized to the average data in the Uniform context (where the timing of the stop signal is unpredictable). Second, the model was fit to the average data in the Early and Late contexts, holding all parameters constant at the best-fitting Uniform values except for one or two parameters of interest. The model that best accounted for differences in the stop-accuracy and RT quantiles across the three context conditions was selected for further investigation of feedback-dependent learning mechanisms. The fitting routine (see Methods section for details) was repeated a total of twenty times using different initialization values for all parameters at the start of each run to avoid biases in the optimization process. The summary of fits to the Uniform Context data is provided in Table 1.

**Table 1.**
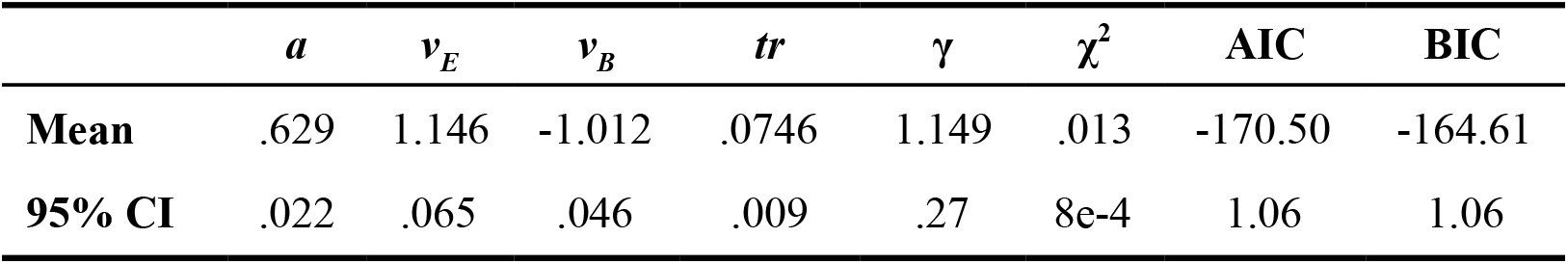
Average Uniform parameters and static DPM fit statistics

We previously found that contextual manipulations of the penalty structure and stop cue probability led to behavioral changes that were best described by modulations in the execution drift-rate when compared to other models with a single free parameter free to vary between conditions [13]. To reduce the combinatorial space of possible model configurations, we first sought to replicate this finding by fitting the model in which only one parameter was free to vary across context condition (see Table 2) - either execution drift (*v*_E_), braking drift (*v*_B_), urgency (γ), or boundary height (*a*). Each model was evaluated using two complexity-penalized goodness-of-fit statistics: Akaike Information Criterion (AIC) and Bayesian Information Criterion (BIC). A difference of 7-10 in the information criteria (IC) values for two models provides strong support for the model with the lower value. In line with our previous findings [13], leaving the execution drift-rate free provided a better account of Context-dependent changes in behavior compared to alternative single-parameter models (Best-Fit AIC*v*_E_ =-369.26; Fig. 5A). The next best fit was afforded by allowing the urgency to vary across conditions (|AIC*v_E_* –AICγ| = 15.83).

**Table 2.**
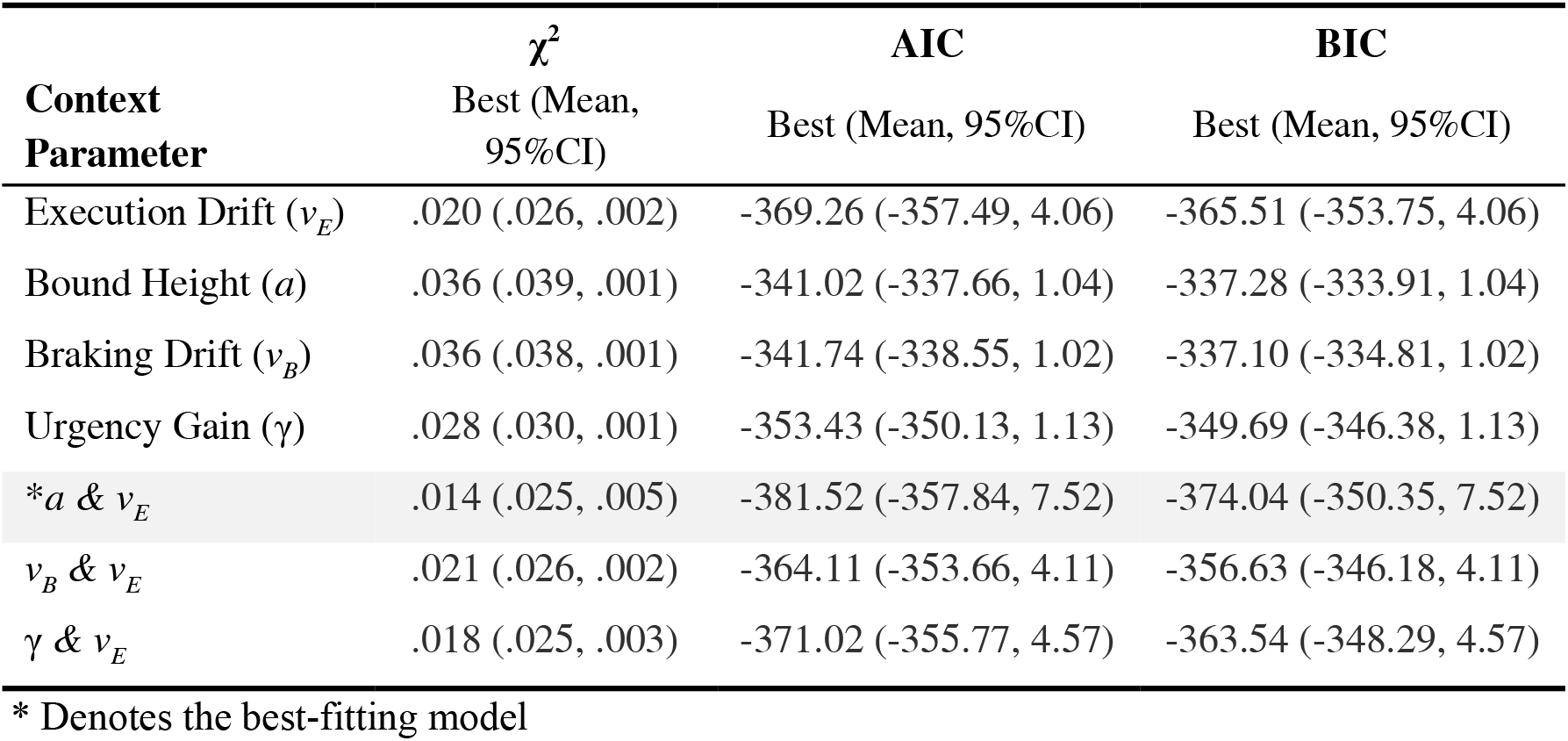
Static Fit Statistics for Early and Late Contexts

**Fig 5.**
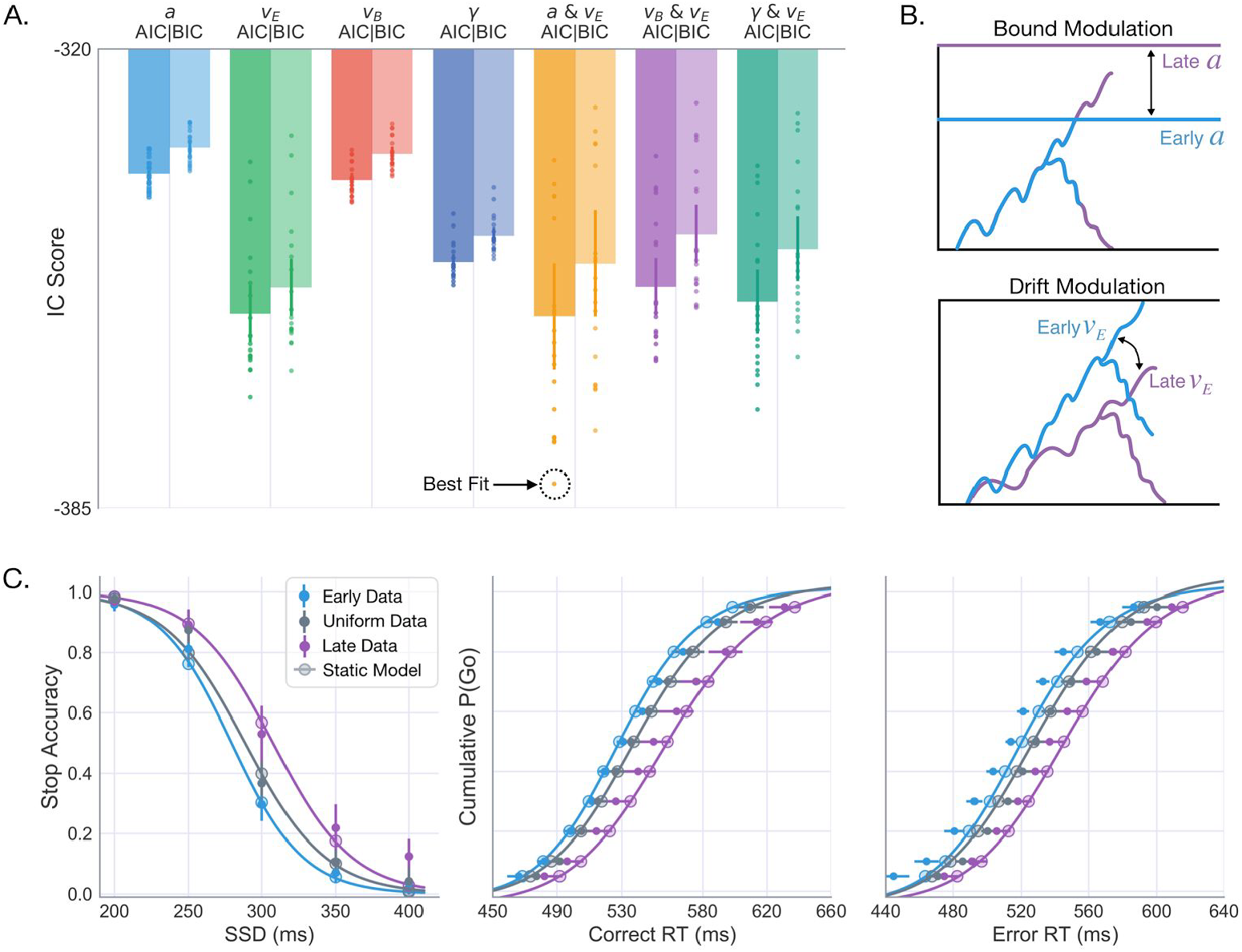
Model comparison and best-fit predictions across context. **(A)** AIC (dark) and BIC (light) scores for all single-parameter models, allowing either execution drift-rate (*v*_*e*_; green), boundary height (*a*; cyan), braking drift-rate (*v*_*b*_; red), or urgency (γ; blue) to vary across Context conditions. Three dual-parameter models were also included to test for possible benefits of allowing *v*_*e*_ (best-fitting single parameter model) to vary along with either *a* (yellow), *v*_*b*_ (purple), or γ (teal). Each dot shows the information criterion (IC) value for one of twenty fits performed with each model. Error bars show the 95% CI. **(B)** Qualitative effects of context on boundary height (top) and drift-rate 14 parameter estimates (bottom) in the Early and Late Contexts. **(C)** Model predicted data (lines and larger transparent circles) simulated with best-fit parameters from the *a* & *v*_*e*_ (corresponding to dotted circle in **A**) overlaid on the average empirical data for Early (blue), Uniform (gray), and Late (purple) context conditions.

To further test the relationship between execution drift-rate and context, we performed another round of fits to test for possible interactions between the execution drift-rate and a second free parameter, either boundary height (*a*), braking drift-rate (*v*_B_), or urgency gain (γ). The AIC and BIC scores from these fits showed that a combination of boundary height and execution drift-rate (*v*_E_ & *a*) provided the best overall fit to the data (Best Fit AIC*a,v*_*e*_ = −381.52), reasonably exceeding that of the drift-only model (|AIC*v*_*E*_ –AIC*γ*| = 12.26) to justify the added complexity of the dual parameter model. Fig 5B shows a qualitative assessment of the *a* & *v*_E_ model’s goodness of fit, revealing a high degree of overlap between the simulated and observed stop-accuracy and RT data in both Early and Late conditions. These results suggest that there may be two targets of learning in the decision process: a strong modulation of the execution drift rate and a more subtle modulation of the boundary height.

### Dual learning mechanisms

It is not clear from the preceding analysis whether error-driven changes in the drift rate and boundary height are able to capture trial-to-trial adjustments of response speed and stop accuracy as statistics of the environment are learned experientially. Here we explore how drift-rate and boundary height mechanisms adapt on a trial-wise basis to different sources of feedback to drive context-dependent control and decision-making.

We implemented two forms of corrective learning - one targeting the execution drift-rate (*v*_E_) and another targeting the height of the decision boundary (*a*). On correct Go trials, the drift-rate parameter was updated according to signed RT errors calculated with respect to the Target RT (T^G^) of 520ms, reducing the drift-rate to slow actions following a “fast” response (Fig. 6A left; RT_*t*_ < 520 ms) and increasing the drift-rate to speed future actions following a “slow” response (Fig. 6A middle, RT_*t*_ > 520 ms). This form of RT-dependent modulation in the drift-rate is motivated by recent findings demonstrating adaptation of action velocity by dopaminergic prediction error signaling in the striatum [25]. In the context of the “believer-skeptic” framework [23], fast RT errors could reinforce the “skeptic” (i.e., indirect pathway) and suppress the “believer” (i.e., direct pathway) by decreasing dopaminergic tone in the striatum.

**Fig 6.**
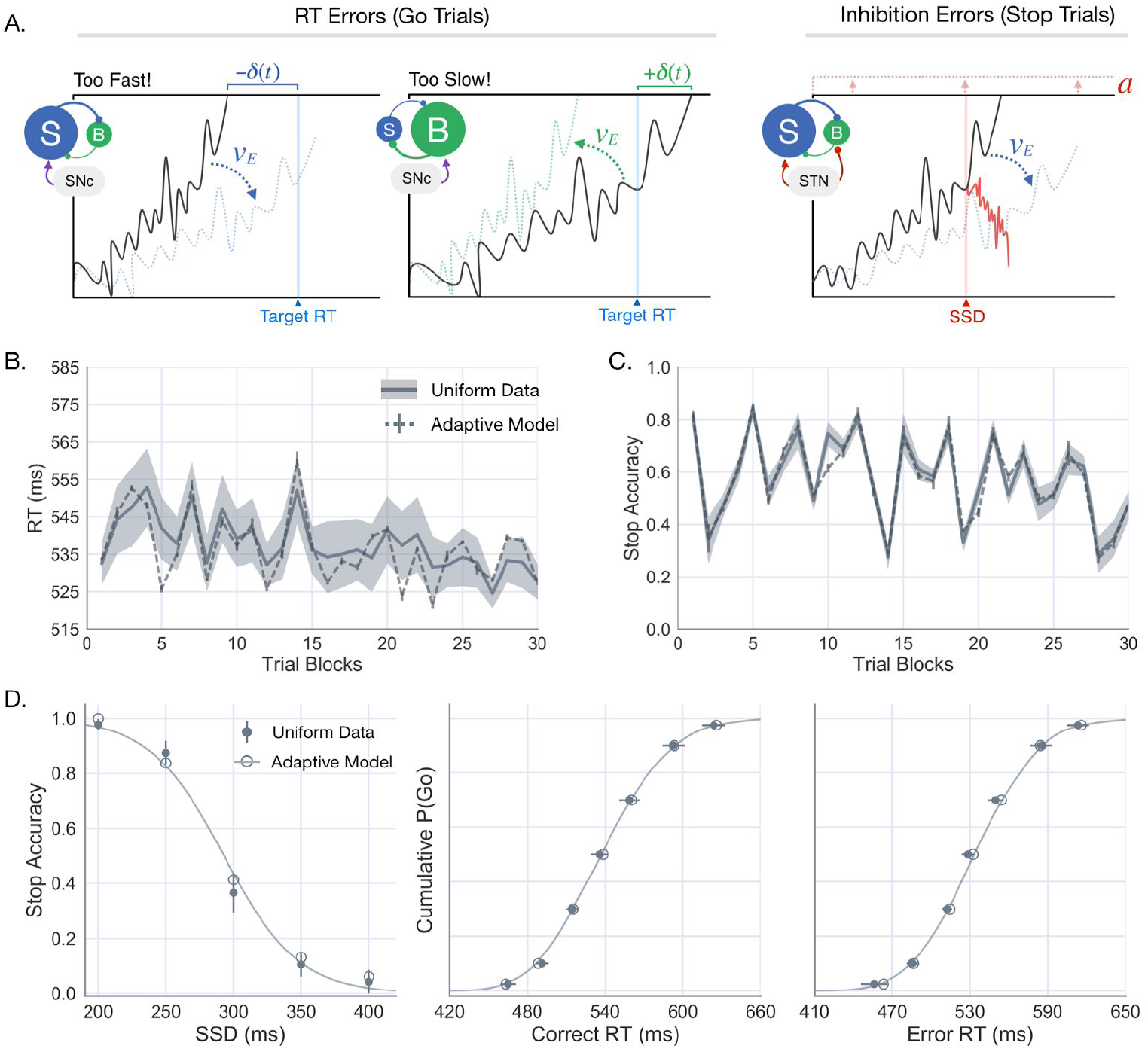
Adaptive DPM and predicted learning trajectory in Uniform condition. **(A)** Schematic showing how the execution drift-rate is modulated following timing errors on Go trials (left) and how the boundary height is modulated following failed inhibitions on Stop trials. **(B)** Subject-averaged timeseries (dark line) and 95% CI (transparent gray) showing the RT on Go trials (left) accuracy percentage on Stop trials (right). Both timeseries are 30 points in total, each calculated as by taking the average RT/stop-accuracy over successive~30-trial windows). The corresponding model predictions are overlaid (light gray line), averaged over simulations to each individual subject’s data. **(C)** Average empirical stop-accuracy and RT statistics in the Uniform condition, (same as shown in Fig 3) with the predictions generated from simulations with the adaptive DPM.

On failed stop trials (*t*_*err*_) the boundary height was increased on the following trial (*t*_*err*_+1) according to a delta function (Fig. 6B) and decayed exponentially on each successive trial until reaching its baseline value or another failed stop was committed (see Methods for details). This form of adaptation in boundary height is motivated by physiological evidence that the STN plays a critical role in setting threshold for action execution and that this relationship is modulated by error commissions [26]. In Fig 6C, the model-predicted change in Go trial RT and Stop trial accuracy across the experiment is overlaid on the corresponding empirical measures, demonstrating a robust model fit to the observed data. To confirm that the trial-averaged behavior of the model was preserved after fitting the learning rates, the stop-accuracy curve and RT statistics were calculated from simulations of the adaptive model and overlaid on the average data from the Uniform condition (Fig 6D). The model’s predictions are indeed closely aligned with all empirical statistics (adaptive χ^2^=.0082, static χ^2^= .013; see Table 3). While this is not necessarily surprising, it is promising to confirm that introducing feedback-dependent adaptation in the drift-rate and boundary height parameters does not compromise the model’s fit to trial averaged statistics.

**Table 3.** Adaptive DPM learning parameters

| $\alpha$ | $\beta$ | $p$ | $\chi^2$ | AIC | BIC |
| --- | --- | --- | --- | --- | --- |
| .24 | .031 | .002 | .0082 | -181.56 | -175.67 |

The fits to the Uniform condition show two possible mechanisms for acquiring the prior on the SSD: adaptive modulation of response speed by the drift-rate and error-driven changes to boundary height. In order to confirm that these mechanisms work together to adaptively learn based only on the statistics of previous input signals, we took the average parameter scheme from the Uniform condition fits and simulated each subject in the Early and Late groups. If the context-dependent changes in the RT distributions and stop-accuracy are indeed a reflection of the proposed learning mechanisms, then the model simulations should reveal similar RT and accuracy time courses as in the observed behavior.

Fig 7A shows the simulated stop-curve and RT distributions generated by the adaptive model based on feedback in the Early and Late conditions. As in the observed data (Fig. 3A), adaptation to Early SSDs led to impaired stopping accuracy, but faster RT’s relative to simulated predictions in the Late condition. In Fig 7B-C, the middle panels show the same trial-binned RT and stop-accuracy means as in Fig 6C (Uniform condition), flanked by corresponding time courses from simulations to Early (left) and Late (right) conditions. The adaptive model predictions show a high degree of flexibility, conforming to idiosyncratic changes in the trial-wise behavioral dynamics within each Context SSD condition. For instance, the RTs in the Early condition exhibit a relatively minor and gradual decay over the course of the experiment (Fig 7B, left), contrasting markedly from the early increase and general volatility of RTs in the Late condition (Fig 7B, right). The adaptive model largely captures both patterns, underscoring feedback-driven adaptation in the drift-rate as a powerful and flexible tool for commanding inhibitory control across a variety of settings. In addition to predicting group differences in the time course of RTs, the simulations in Fig 7C show a striking degree of precision in the model-estimated changes in stop-accuracy, both over time and between groups.

**Fig 7.**
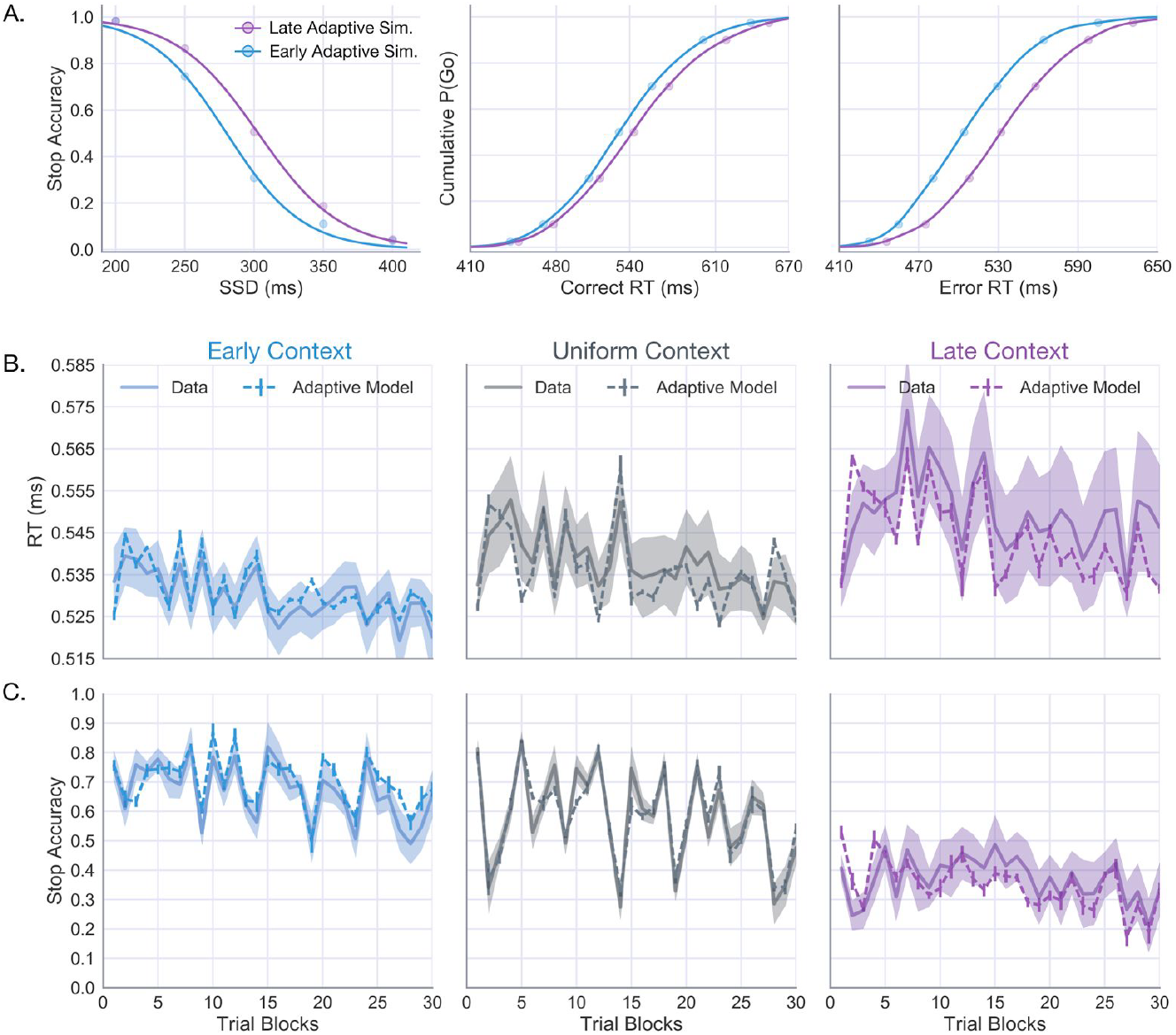
Adaptive DPM modulates behavior to context-specific control demands. **(A)** Average stop-accuracy curves (left) and correct (middle) and error (right) RT distributions predicted by adaptive model simulations in the Early (blue) and Late (purple) Contexts (initialized with the optimal parameters of the Uniform Context). **(B)** Empirical timeseries of Go RT’s with model predictions overlaid (calculated using the same method described for Fig 5B-C for Early (left), Uniform (middle, same as in Fig 5C), and Late (right) Contexts. **(C)** Empirical and model predicted timeseries of stop-accuracy for the same conditions as in **B**.

Because the static model fits revealed marginal evidence for the drift-only model (Fig. 5A), we next asked whether this simpler model was able to account for the learning-related behavioral changes (see Fig. 6B) with the same precision as the dual learning (i.e., drift and boundary) model. To test this hypothesis, we ran simulations in which the boundary learning rate was set to zero, thereby leaving only the drift-rate free to vary in response to feedback. Fig 8A shows the error between observed and model-predicted estimates for each of the behavioral measures in Fig 2 (e.g., RT, stop accuracy, and post-error slowing) based on 25 simulations of the drift-only and dual learning models. Compared to the drift-only model, the dual learning model showed no significant benefits in terms of fit to the trial-wise RT, *t*(24)=1.09, *p*=.28, or accuracy *t*(24)=.23, *p*=.82, but showed a marked improvement in the fit to post-error slowing, *t*(24)=-6.91, p<.00001 (Fig. 8A).

**Fig 8.**
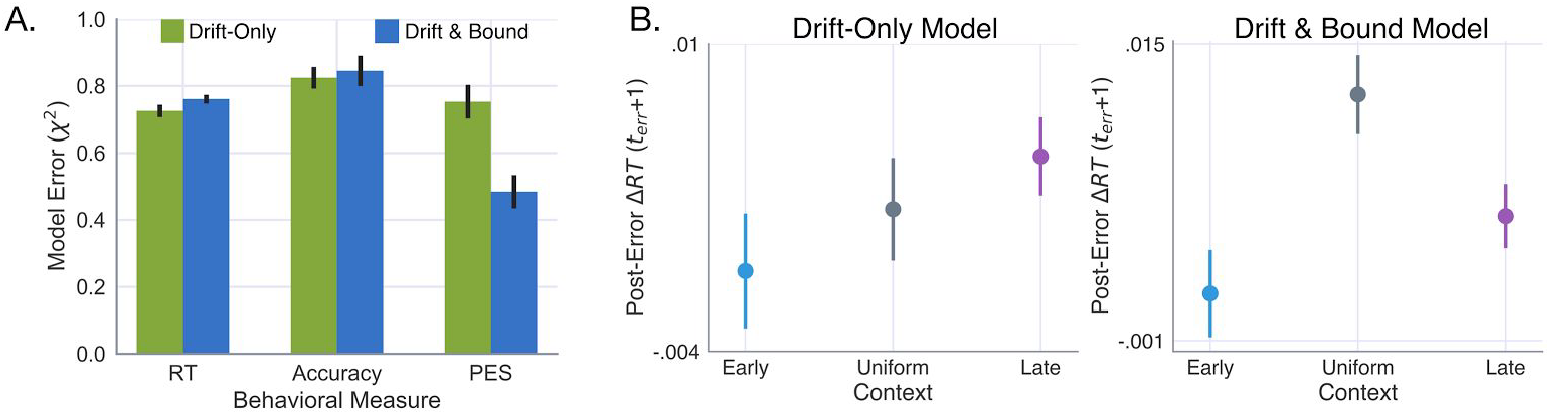
Predictive utility of including boundary adaptation compared to drift-only model. (**A**) Relative error of simulated compared to observed RT, accuracy, and post-error slowing measures based on twenty simulated data sets for the drift-only and drift & bound adaptive models. Post-error slowing in each context condition as predicted by the **(B)** drift-only and **(C)** drift & bound models. Error bars reflect the 95% CI around the mean.

Importantly, the interaction of drift-rate and boundary adaptation in the dual learning model not only reduced the error in the model fit, but recovered the same qualitative pattern of post-error slowing across Contexts observed in the data (Fig. 8B). In contrast, the drift-only model predicted the largest post-error slowing effect in the Early condition (Fig. 8B, left). This is particularly revealing since no information about the observed post-error slowing was included in the adaptive cost function when fitting the learning-rate parameters. Collectively, these results suggest that goal-directed tuning of movement timing (RT) and control (stop accuracy) is best described by feedback-driven changes in the drift-rate and boundary-height parameters of accumulation-to-bound decisions.

## Discussion

The current results demonstrate the existence of two separable, yet interacting, learning mechanisms for adapting inhibitory control. Adaptation to errors in the timing of action execution was mediated by adjustments in the drift rate that progressively improved the precision of RTs with respect to the target response time. Inhibition errors, i.e., executed responses on trials requiring a stop, had a post-error slowing effect, mediated by an increase in the execution threshold that decayed over subsequent trials. These two mechanisms allowed for principled, context-specific adjustments in behavioral control to conflicting sources of task error, i.e., go timing and stop accuracy. Relative to the Uniform condition, subjects in the Early condition exhibited faster RTs at the expense of accuracy on probe Stop trials (see Fig. 3B). Subjects in the Early condition benefited from predictably short SSDs, making it easier to reactively cancel actions on Stop trials without sacrificing the the precision of action timing on Go trials. Over time, these subjects experience relatively few stop errors and thus focused on minimizing errors between their Go trial RT and the target response times (see Fig. 6B, left). In contrast, subjects in the Late condition slowed their RT on Go trials to accommodate the higher probability of a stop-cue late in the trial, introducing greater long term costs due to more frequent stop errors, and thus, delay responding on Go trials to improve inhibition accuracy. This principled adaptation of control is only possible due to the presence of dual learning mechanisms on the action control algorithm.

### Dual learning mechanisms for adaptive control

In accumulation-to-bound models of decision-making, the drift-rate and boundary height parameters are functionally dissociated as representing the strength of evidence and response caution, respectively [4,37]. However, this dissociation is only useful insofar as evidence is clearly defined and can be manipulated independently of the additional factors bearing on behavior (e.g. caution, expectation, attention). With the exception of perceptual decision-making tasks [38], where evidence can be interpreted with respect to the strength of sensory information provided by the stimulus, it is often unclear what sources of information are actually encoded as evidence in the decision process and what are used to set the boundary height. In the following section we discuss evidence that, within the context of go/no-go control tasks, drift-rate and boundary height mechanisms are driven by corticostriatal and corticosubthalamic systems, respectively; and furthermore, that both participate in feedback-dependent learning but in the service of distinct behavioral goals.

### Drift-rate: dopaminergic modulation of corticostriatal pathways

At the computational level, the drift-rate parameter reflects the log-likelihood ratio of evidence for alternative hypotheses. In the context of the current task, the execution drift-rate can be interpreted as representing the relative evidence for go and no-go decisions encoded by the circuit-level competition between the direct and indirect pathways [39]. Indeed, studies combining behavioral modeling with single-unit recordings [16], optogenetics in animals [25], and neuroimaging in humans [40] have found reliable links between behaviorally derived estimates of drift-rate and activity in the striatum [28]. Crucially, the dynamics of competition between direct and indirect pathways is sensitive to dopaminergic signals that provide important feedback about the environmental consequences of recent actions to drive behavior in the direction of the agent’s current goal [41–44]. Feedback-dependent reweighting of corticostriatal connections has primarily been studied in the context of action-value learning; however, new evidence suggests a more nuanced role in tuning task-relevant movement parameters [25,45,46]. Yttri and Duman [25] demonstrated this by stimulating direct or indirect pathway neurons in the mouse striatum based on the velocity of a recently executed lever press and measuring the effects on future movements. Similar to the opponent effects of dopaminergic error signals that mediate action-value associations [41,47], they found that stimulation of the direct pathway following high-velocity presses further increased the velocity of future movements whereas stimulation of indirect pathway neurons decreased velocity. While the current study was not concerned with action velocity, per se, the adaptation of the drift-rate parameter to errors in action timing resembles a similar behavioral dynamic to that observed by Yttri and Dudman [25]. Indeed, a recent study by Soares et al., [48] found that dopaminergic neurons in the mouse midbrain were not only necessary for accurate temporal perception, but that the perception of time could be systematically sped up or slowed down through optogenetic suppression and stimulation of these neurons. Future studies will be needed to confirm the proposed dependency of the drift-rate on striatum in which model-fits to behavior are performed in the presence of dopaminergic weighting at direct and indirect synapses.

Based on previous evidence that proactive control is mediated by the striatum, we have argued that the feedback-dependent modulation of execution drift-rate from to changes in the stop-cue prior is, at least in part, a reflection of the competition between the direct and indirect pathways, and that the observed adaptation in this parameter is a reflection of dopaminergic feedback about errors in action timing. In addition to the dopamine hypothesis, an alternative possibility is that adaptation the drift-rate stems from top-down changes in the background excitability of the striatum, driven by diffuse inputs from premotor regions such as supplementary motor area (SMA) and pre-SMA [18,40,49,50]. It remains unclear what functional differences may exist between premotor and dopaminergic representations of time or how they might differentially influence the encoding of action timing within the striatum. Integrating the behavioral and modeling techniques defined here with electrophysiological and optogenetic manipulations can better distinguish the nature of the training signal that modulates striatal activity during action control.

### Boundary height: post-error slowing by the STN

The striatum has long been the focus of investigations into the neural basis of feedback-dependent learning; however, BG pathways have multiple targets of plasticity beyond the striatum. Recently, similar links have been identified between activity fluctuations in the STN and adaptive changes in behavior [17,26,51,52], raising new and interesting questions about the extent to which striatal and subthalamic learning signals independently influence behavior and how they might interact [53]. Numerous studies have implicated the STN in setting the height of the decision threshold [9,17,29,54,55], controlled by diffuse excitatory inputs to the output nucleus of the BG and further suppressing motor thalamus to delay action execution (see Fig. 1A). Due to the monosynaptic connections between cortex and the STN that make up the hyperdirect pathway [56], unexpected sensory events (e.g., stop signals) can be quickly relayed through the STN to raise the decision threshold for ongoing action plans to prevent execution [57]. In addition to this rapid cortically-mediated form of adaptation, evidence suggests that strategic adjustments in decision threshold are achieved via activity-dependent plasticity in the connections between STN and GPe [24,58]. In the current study, adaptive changes in the boundary height accounted for the observed post-error slowing in responses following failed stop trials, motivated by neuroimaging and electrophysiological evidence STN-mediated slowing of responses [9,17,26]. For simplicity, boundary adaption was restricted to being unidirectional - increasing after a stop-error and decaying back to, but never below, its original value. However, some evidence suggests that STN exerts bidirectional control over decision threshold, capable of promoting the adoption of both speed and accuracy policies [29]. Thus, relating the adaptive threshold in the dependent process model to recordings in the STN will likely require a more nuanced approach in order to generalize beyond the current task. Future studies will be needed to examine how each of these neural mechanisms are recruited to modify behavior, the relevant contexts and task dimensions they are sensitive to and the timescales they operate on.

### Conclusion

Computational modeling of RT and accuracy in an adaptive stop-signal task revealed two independent learning mechanisms underlying feedback-dependent changes in control - one responsible for gradually tuning the execution drift-rate to timing errors on Go trials and another for increasing caution after a failed Stop trial by raising the execution boundary. The tuning of the drift-rate and boundary height parameters supports recent evidence of striatal-and STN-driven changes in goal-directed behavior following errors in action timing and inhibitory control respectively. While cognitive modeling approaches are unable to capture the complexity of neural information processing that underlies adaptive action control, they provide a rich description of the component operations and can thus be exploited for the purpose of more keenly parsing the functional role of different neural substrates. The current study shows how a straightforward hybridization of two cognitive modeling frameworks - accumulation-to-bound dynamics with basic principles of reinforcement learning - provides confirmatory evidence for a dual-mechanism account of feedback dependent learning in inhibitory control.

## Methods

### Participants

Neurologically healthy adult participants (N=75, Mean age 22 years) were recruited from the Psychology Research Experiment System at Carnegie Mellon University and compensated for their participation through course credit toward fulfillment of their semester course requirements. All experimental and analytical protocols described in this study were approved by the local Institutional Review Board (IRB) at Carnegie Mellon University. Experimenters obtained informed, written consent from all subjects in compliance with IRB guidelines.

### Adaptive stop-signal task

All subjects completed a stop-signal task (N_trials_=880) in which a vertically moving bar approached a white horizontal target line at the top of the screen (Fig. 2A). On ‘Go’ trials (N_GO_=600) the subject was instructed to make a key press as soon as the bar crossed the target. The bar always intersected the target line at 520ms after trial onset. On each trial, the bar continued filling upward until a keypress was registered or until reaching the top of the screen, allowing a 680ms window for the subject to make a response. If no response was registered the subject received a penalty of (-100pts). On Go trials where a response was recorded before the 680 ms trial deadline, the subject received a score reflecting the precision of their response time relative to the target intersection, resulting in maximal points when RT=520ms. On Stop trials, the bar would stop and turn red prior intersecting the target line, prompting the subject to withhold their response. Successful and unsuccessful Stop trials yielded a reward of +200 points and penalty of −100 points, respectively. On the majority of Stop trials, the stop-signal delay (SSD) - the delay between trial onset and when the bar stopped – was sampled from a specific probability distribution (see Fig. 2B). We refer to these trials as Context Stop trials (N_Context_=200). Context SSD’s in the Early and Late groups were sampled from Gaussian distributions with equal variance (σ=35ms), centered at μ_E_=250ms and μ_L_=350ms, respectively. Context SSDs in the Uniform group were sampled from a uniform distribution spanning a 10-520ms window. In Fig 2B, the sampled SSD times are plotted for a single subject in each Context – shown as dashes on a timeline ranging from 0-520ms. Finally, additional probe Stop trials (N_Probe_=80) were included in which the bar stopped at 200, 250, 300, 350, or 400 ms after trial onset (16 trials per probe SSD), shown at the bottom of the Fig 2B timeline as red dashes.

## Computational Models

### Dependent Process Model

The dependent process model (Fig. 1C; [13]) assumes that action-facilitating (i.e., direct) and action-suppressing (i.e., indirect) signals are integrated over time as a single execution process (θ_E_), with a drift-rate that increases with the ratio of direct-to-indirect pathway activation. The linear drift and diffusion (φ_E_) of the execution process is described by the stochastic differential equation in equation 1, accumulating with a mean rate of *v*_E_ (i.e., drift rate) and a standard deviation described by the dynamics of a white noise process (*dW*) with diffusion constant σ. The execution process is fully described by equation 2 in which the linear accumulation described by equation 1 is scaled by an urgency signal, modeled as a hyperbolic cosine function of time with gain γ.

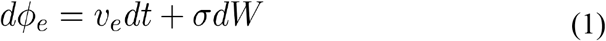

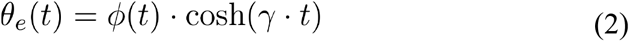

A response is recorded if θ_E_ reaches the execution boundary (*a*) before the end of the trial window (680ms) and before the braking process reaches the lower (0) boundary (see below). In the event of a stop cue, the braking process (θ_B_)is initiated at the current state of θ_E_ with a negative drift rate (*v*_B_). If θ_B_ reaches the 0 boundary before θ_E_ reaches the execution boundary then no response or RT is recorded from the model. The in θ_B_ over time is given by equation 3, expressing the same temporal dynamics of φ_E_ but with a negative drift rate. The dependency between θ_B_ and θ_E_ in the model is described by the conditional statement in equation 4, declaring that the initial state of θ_B_ (occurring at t = SSD) is equal to the state of θ_E_(SSD).

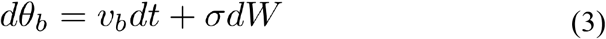

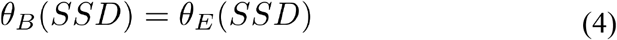

In order to determine which of the model parameter(s) best accounted for the observed behavioral effects across Contexts, we first fit the model to the average data in the Uniform group, leaving all parameters free (see Table 1). Using the optimized Uniform parameter estimates to initialize the model, we then fit different versions of the model to data in the Early and Late groups allowing only one or two select parameters to vary between conditions. This form of model comparison provides a straightforward means of testing alternative hypotheses about the mechanism underlying Context-specific adaptation. The fitting procedure utilized a combination of global and local optimization techniques [13,59]. All fits were initialized from multiple starting values in steps to avoid biasing model selection to unfair advantages in the initial settings. Given a set of initial parameter values, all model parameters – execution drift-rate (*v*_E_), braking drift-rate (*v*_B_), execution onset delay (*tr*), boundary height (*a*) and dynamic gain (γ) - were optimized by minimizing a weighted cost function χ_*static*_ (see eq. 5) equal to the summed and squared error between an observed and simulated (denoted by *^*) vector containing the following statistics: probability (*P*)of responding on Go trials (*g*), probability of stopping at each Probe SSD (*d*={200, 250, 300, 350, 400ms}), and RT quantiles (q={.1, .2,…, .9}) on correct (RT^C^) and error (RT^E^) trials.

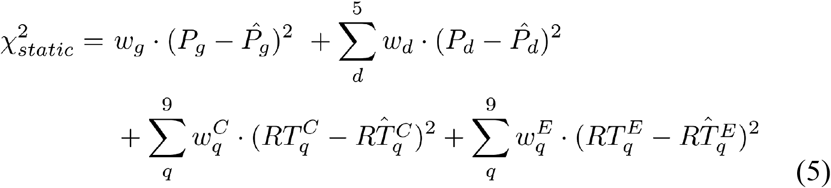

The cost-function weights (ω) were derived by first taking the variance of each summary measure included in the observed vector (across subjects), then dividing the mean variance by the full vector of variance scores. This approach represents the variability of each value in the vector as a ratio [60], where values closer to the mean are assigned a weight close to 1, and values associated with higher variability a weight <1, lower variability a weight >1 [13,61]. Weights applied to the RT quantiles were calculated by estimating the variance for each of the RT quantiles [62] and then dividing the mean variance by that of each quantile. Stop accuracy weights were calculated by taking the variance in stop accuracy at each Probe SSD (across subjects) and then dividing the mean variance by that of each condition.

In order to obtain an estimate of fit reliability for each model we restarted the fitting procedure from 20 randomly sampled sets of initial parameter values. Each initial set was then optimized to average data in the Uniform condition using the basinhopping algorithm [63] to find the region of global minimum followed by a Nelder-Mead simplex optimization [64] for fine tuning globally optimized parameter values. The simplex-optimized parameter estimates were then held constant except for one or two designated context-dependent parameter(s) that were submitted to a second Simplex run in order to find the best fitting values in the Early and Late conditions.

### Parameter recovery of static DPM

Parameters were initially sampled from the following distributions:

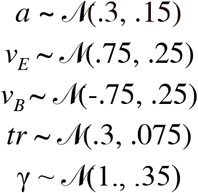

where 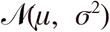 represents a normal distribution with mean *μ* and standard deviation σ. A total of 2000 parameter sets were initially sampled and used to simulate vectors of stopping accuracy and RT quantiles that were compared to those of the average subject in each Context by means of a equation 5. For each of the three Context conditions, the corresponding sampled set of parameters associated with the lowest error value was then selected as a ‘group-level’ parameter set. For each of the three ‘group-level’ parameter sets, 20 datasets were generated by sampling 25 ‘subject-level’ parameter sets that were then used to simulate 1000 trials each. Subject-level parameters for each of the three groups were sampled from normal distributions with means and standard deviations shown in Table 4.

**Table 4.** Means and standard deviations of subject-level parameter samples

| **True Sets** | $\mu_a$<br>($\sigma=.01$) | $\mu_{vE}$<br>($\sigma=.01$) | $\mu_{vB}$<br>($\sigma=.01$) | $\mu_{tr}$<br>($\sigma=.005$) | $\mu_\gamma$<br>($\sigma=.02$) |
| --- | --- | --- | --- | --- | --- |
| **Set 1** | .358 | .946 | -.410 | .168 | .926 |
| **Set 2** | .367 | .882 | -.424 | .169 | 1.152 |
| **Set 2** | .322 | .789 | -.392 | .183 | 1.155 |

### Adaptive DPM

On correct Go trials, the drift-rate (*v*_*t*_) was updated according to equation 6 to reflect the signed difference between model’s response time on the current trial and the Target time (T^G^ = 520ms), increasing the drift-rate following “slow” responses (i.e., RT_t_>T^G^, Fig. 5A left) and decreasing the drift-rate following “fast” responses (i.e., RT_t_<T^G^, Fig. 5A middle). On failed stop trials, *v_t_* was updated according to the same equation but with the error term reflecting the difference between RT_t_ and the trial response deadline (T^S^=680), thus, slowing the drift-rate to reduce the probability of failed stops in the future.

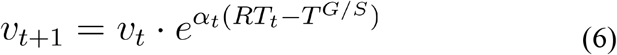

The boundary height (*a*_0_) adapted to failed stop trials (Fig. 5A right) by increasing according to a delta function with height β_t_ and decaying exponentially on each subsequent trial (*a_terr_*) until reaching its baseline value *a*_0_ or until another stop error occurred (eq. 7).

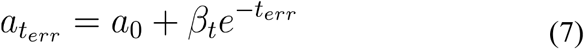

On all correct Go trials and the first failed stop trial, the timing errors were scaled by the same learning rate (α_0_). An additional parameter was included to modulate the sensitivity (*π*) to stop errors over time, calculated according the power rule shown in equation 8. The trial-wise stop-error sensitivity was used to scale α_t_ (eq. 9) and β_t_ (eq. 10) learning rates on failed stop trials before calculating the drift (eq. 6) and boundary height (eq. 7) updates.

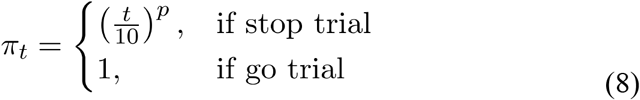

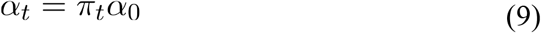

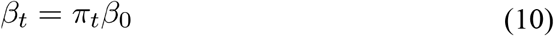

To obtain estimates for the learning rate parameters for adaptation in the drift-rate (α), boundary-height (β), and sensitivity to stop errors (*p*) this adaptive form of the model was re-fit to the RT and stop accuracy data in the Uniform condition, holding all previously estimated parameters constant. Because standard parameter optimization for accumulator models requires information about the variance of response-times across trials, these approaches are poorly suited for investigating how decision parameters respond to error on a trial-wise basis. To overcome this issue, cost function was modified (*χ*_*adapt*_) to identify the values for α, β, and *p* that minimized the sum of the weighted difference between the average observed and model-predicted stop accuracy (*μ*_*acc*_) and Go RT (*μ*_*rt*_) over a moving window of roughly 30 trials (30 bins total; eq. 11). The weights applied to the model-predicted error in stop accuracy (*ω*_*acc*_) and RT (*ω*_*rt*_) were calculated using the same method as for the static model cost function, assigning less weight to estimates in bins (*i*) with higher observed variance across subjects. By averaging the behavioral measures in 30-trial bins, this ensured that multiple Stop trials were included in each bin while still allowing relatively high-frequency behavioral changes to be expressed in the cost function.

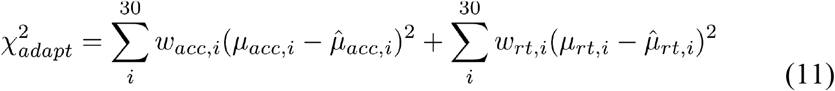

These fits were performed by iteratively simulating the same trial sequence as observed for each individual subject, and fitting the average simulated subject to the average observed subject. This ensures that direct comparisons can be made between the trajectory of learning in the model and actual behavior.

## Acknowledgments

The authors would like to thank Patrick Beukema and Kevin Jarbo for their helpful comments on early drafts of this manuscript and Leo Scholl and Tara Molesworth for assistance with data collection.

